# Deletion of immune evasion genes provides an effective vaccine design for tumor-associated herpesviruses

**DOI:** 10.1101/2020.04.13.034082

**Authors:** Gurpreet Brar, Nisar A. Farhat, Alisa Sukhina, Alex K. Lam, Yong Hoon Kim, Tiffany Hsu, Leming Tong, Wai Wai Lin, Carl F. Ware, Marcia A. Blackman, Ren Sun, Ting-Ting Wu

**Author notes:** Corresponding author: Ting-Ting Wu. These authors contributed equally to this work (Gurpreet Brar, Nisar A. Farhat).

## Abstract

Vaccines based on live attenuated viruses often induce broad, multifaceted immune responses. However, they also usually sacrifice immunogenicity for attenuation. It is particularly difficult to elicit an effective vaccine for herpesviruses due to an armament of immune evasion genes and a latent phase. Here, to overcome the limitation of attenuation, we developed a rational herpesvirus vaccine in which viral immune evasion genes were deleted to enhance immunogenicity while also attaining safety. To test this vaccine strategy, we utilized murine gammaherpesvirus-68 (MHV-68) as a proof-of-concept model for the cancer-associated human γ-herpesviruses, Epstein-Barr virus and Kaposi sarcoma-associated herpesvirus. We engineered a recombinant MHV-68 virus by targeted inactivation of viral antagonists of type I interferon (IFN-I) pathway and deletion of the latency locus responsible for persistent infection. This recombinant virus is highly attenuated with no measurable capacity for replication, latency, or persistence in immunocompetent hosts. It stimulates robust innate immunity, differentiates virus-specific memory T cells, and elicits neutralizing antibodies. A single vaccination affords durable protection that blocks the establishment of latency following challenge with the wild type MHV-68 for at least six months post-vaccination. These results provide a novel approach to effective vaccination against cancer-associated herpesviruses through the elimination of latency and key immune evasion mechanisms from the pathogen.

## Introduction

Human γ-herpesviruses Epstein-Barr virus (EBV) and Kaposi sarcoma-associated herpesvirus (KSHV) are associated with cancer, and with no effective vaccine remain a global health challenge. Despite strong innate and adaptive immune responses, once acquired, herpesviruses persist for the rest of the host’s life. EBV is associated with Burkitt’s lymphoma, nasopharyngeal carcinoma (NPC), and Hodgkin’s- and non-Hodgkin’s lymphomas^1–3^ while KSHV is associated with Kaposi’s sarcoma (KS), primary effusion lymphoma (PEL), and multicentric Castleman’s disease (MCD). These malignancies frequently develop in AIDS patients^4–6^, but also in immunocompetent people with more than 160,000 annual cancer cases associated with EBV and KSHV^7^. Clearly, effective vaccines against human γ-herpesviruses would dramatically reduce the incidence of malignancies associated with these viruses.

Herpesviruses establish persistent infections characterized by lytic replication and latency. Lytic replication of α- and β-herpesviruses results in disease pathologies, such as varicella and herpes zoster for Varicella-Zoster virus (VZV), cold sores and genital lesions for herpes simplex virus (HSV), and congenital defects for cytomegalovirus (CMV). In comparison, malignancies associated with γ-herpesvirus infection are linked to viral latency. Viral genes expressed during latency promote the survival and proliferation of infected cells with increased susceptibility to carcinogenic transformation. Therefore, effective vaccine strategies against tumor-associated herpesviruses ideally should prevent latent infections. The oncogenic potential of γ-herpesviruses has focused vaccine research and development on protein subunit vaccines without the latency risk of live viruses. Subunit anti-EBV vaccines have been based on the envelope protein gp350. Antibodies against gp350 block EBV infection in B-cells^8^ which are long-term latency reservoirs. Gp350-based vaccines protect against infectious mononucleosis (IM); however, they do not influence the overall infection rate^9^ and thus are unlikely to prevent EBV-associated cancers. Similarly, subunit vaccines against HSV-2 may reduce genital lesions but do not prevent infection^10^. Therefore, a new strategy is required to establish wide, durable immunity against herpesviruses.

Live viral vaccines simulate an infection presenting the entire viral antigen repertoire to create stable, long lasting immune memory. Viruses can be attenuated by removing viral genes essential for replication. However, replication-deficient viruses may undergo recombination and regain replication capacity during propagation in complementing cells expressing the missing genes. Furthermore, attenuation of replication competence may compromise immunogenicity. An alternative approach is to selectively inactivate viral genes involved in immune evasion in order to attenuate replication and enhance immunogenicity. Viral antagonists of type I interferon (IFN-I) response are an important class of immune evasion genes to consider. The IFN-I response is the first line of antiviral defense in the host, and subverting the IFN-I response is critical for viruses to establish infections in hosts. IFN-I response initiates a signaling cascade inducing the transcription of over 300 genes that counteract viral infections^11–14^ and also promotes adaptive immune responses. Approximately 25% of genes encoded by γ-herpesviruses modulate host immunity, including those that counteract the IFN-I response^15,16^.

Here, we designed a viral vaccine that addresses both immunogenicity and safety. We hypothesized that a recombinant herpesvirus lacking multiple IFN-I evasion genes and deficient in latency can prime memory development in T and B cells despite attenuated replication. However, human γ-herpesviruses are highly species-specific and cannot infect small animals. To overcome this limitation, we utilized murine gammaherpesvirus 68 (MHV-68), closely related to EBV and KSHV^17^, to test the hypothesis. We show that an MHV-68 virus engineered to be latency- and immune evasion-deficient is highly attenuated in immunocompetent hosts yet a potent inducer of antiviral immunity. Moreover, this recombinant virus elicits robust long-lasting protection against persistent wild type viral infection.

## Results

### Construction of a virus deficient in immune evasion and persistence (DIP)

In our previous genome-wide screen of MHV-68 open reading frames (ORFs), eight were found to reduce IFN-I responses according to an IFN-stimulated response element (ISRE) reporter assay^18^. We selected *ORF10, ORF36*, and *ORF54* for the insertion of translational stop codons as these genes are dispensable for viral replication and are conserved among MHV-68, KSHV, and EBV. We also inactivated K3, a viral inhibitor of MHC class I antigen presentation pathway, by truncation to increase the immunogenicity of the vaccine virus^19,20^. We hypothesized that removal of these four immune evasion genes would increase immunogenicity while attenuating replication of the vaccine virus by inducing a robust IFN response and presenting all viral epitopes.

A critical safety component of our design is eliminating the latency of the vaccine virus. In KSHV and MHV-68, the biphasic life cycle is regulated by RTA, the replication and transcription activator, and by LANA, the latency associated nuclear antigen. The latter is required for latency establishment^21–23^ while the former upregulates lytic genes^24–26^. We previously showed that abolishing LANA expression combined with constitutive RTA expression results in a latency-deficient virus^27^. Here, we replaced the latency locus comprising ORF72, ORF73 (LANA), ORF74, and M11 with constitutively expressed RTA driven by the phosphoglycerate kinase 1 (PGK) promoter in a two-tiered approach to prevent persistent infection. Deletion of the latency locus, constitutive RTA expression, and the removal of immune evasion genes created a live attenuated γ-herpesvirus vaccine named DIP (<u>d</u>eficient in immune evasion and <u>p</u>ersistence) (Fig. 1A).

**Figure 1.**
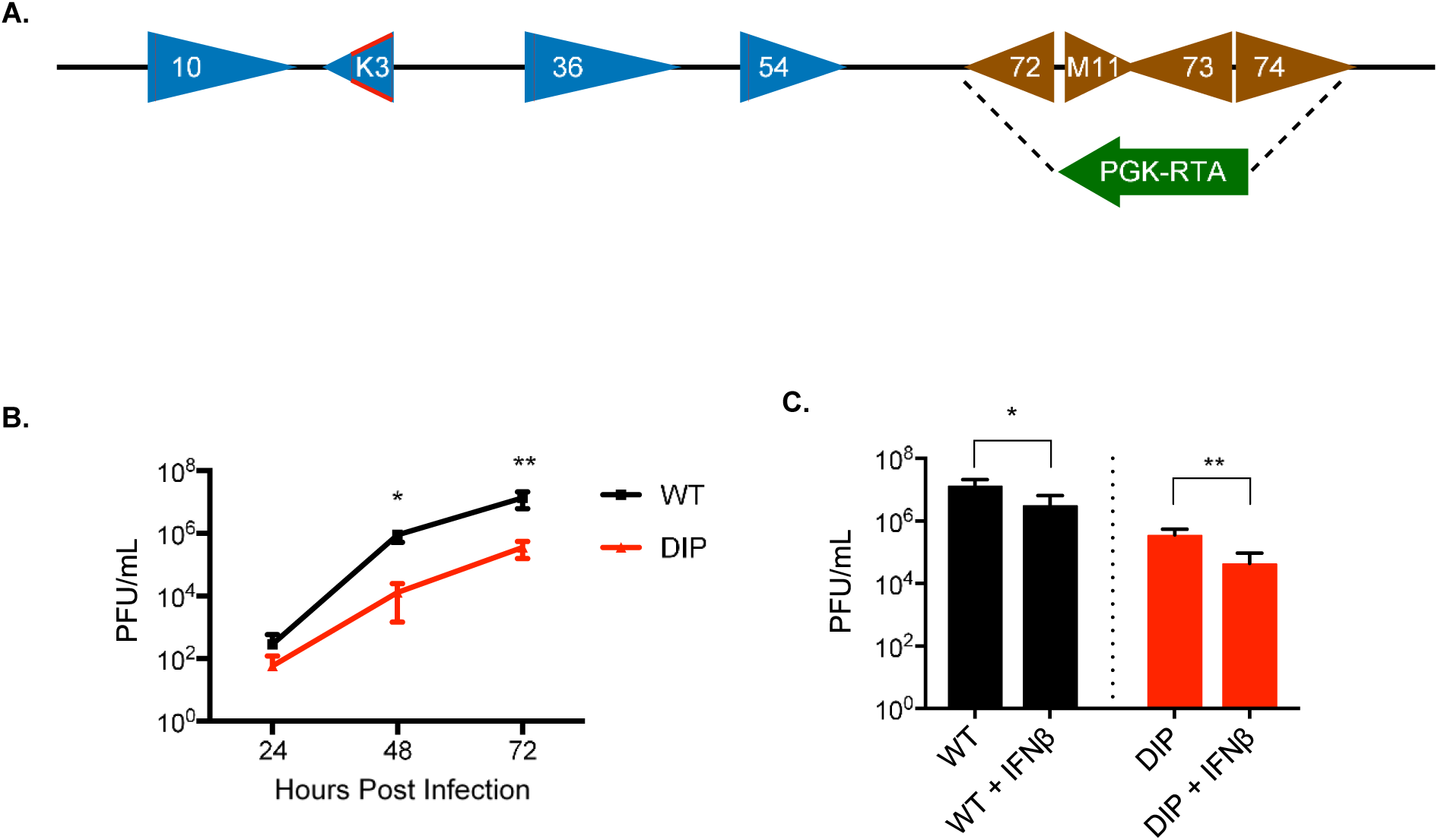
Construction of DIP virus and its replication properties in vitro. (**A**) Schematic representation of mutations introduced in the MHV-68 genome to generate the DIP vaccine. Red lines indicate insertion of translation stop codons into ORF10, ORF36, and ORF54. The open red tetragon indicates deletion of the coding sequence in K3. The latency locus was replaced by the RTA cassette (arrowhead) constitutively driven by the PGK promoter. (**B**) Growth curves of the WT and DIP viruses in 3T3 cells using MOI = 0.01 and measured by plaque assay to quantify virion production. (**C**) NIH 3T3 cells were either mock treated or treated with 100 U mL^-1^ IFN-β for 24 h then infected with either WT or DIP virus at MOI = 0.01 for 72 h. Virion production was quantified with plaque assays. All experiments were performed in triplicate and statistical significance was analyzed by a two-tailed Student’s *t*-test. Graphs represent means of triplicates with SD.

### DIP replication is attenuated in vitro

Comparison of the *in vitro* growth kinetics of DIP in NIH3T3 fibroblasts with the wild type (WT) virus showed that DIP replication was significantly attenuated. After infection at MOI of 0.01, DIP yielded 300-fold and 40-fold less viral production than WT at 48 h and 72 h post-infection, respectively (Fig. 1B). Pretreatment with IFN-β inhibited replication of WT by 10-fold and DIP by 100-fold (Fig. 1C). This larger decrease in infectious DIP virion production confirmed augmented susceptibility to the IFN-I response in the absence of viral IFN evasion genes.

### DIP produces no infectious virions in vivo

We hypothesized that removal of the viral IFN-I evasion genes would generate a highly attenuated vaccine *in vivo*. To test this, we infected C57Bl/6 mice intraperitoneally and harvested their spleens 3 d after infection. While the WT virus produced 88 PFU/spleen, no infectious virus was detected in the spleens of DIP-inoculated mice (Fig. 2A). We also harvested spleens at later times post-infection and assessed spontaneously reactivating virus by the infectious center assay and quantified latent viral genomes by qPCR. Viral reactivation or latent virus was undetectable in the spleens of DIP-infected mice at 14 d (Figs. 2B and 2C) and at 2 mo (Figs. 2E and 2F) after infection. Furthermore, no infectious virion production in the lungs or latency establishment was observed in the spleens after intranasal inoculation (Figs. S1A-D).

**Figure 2.**
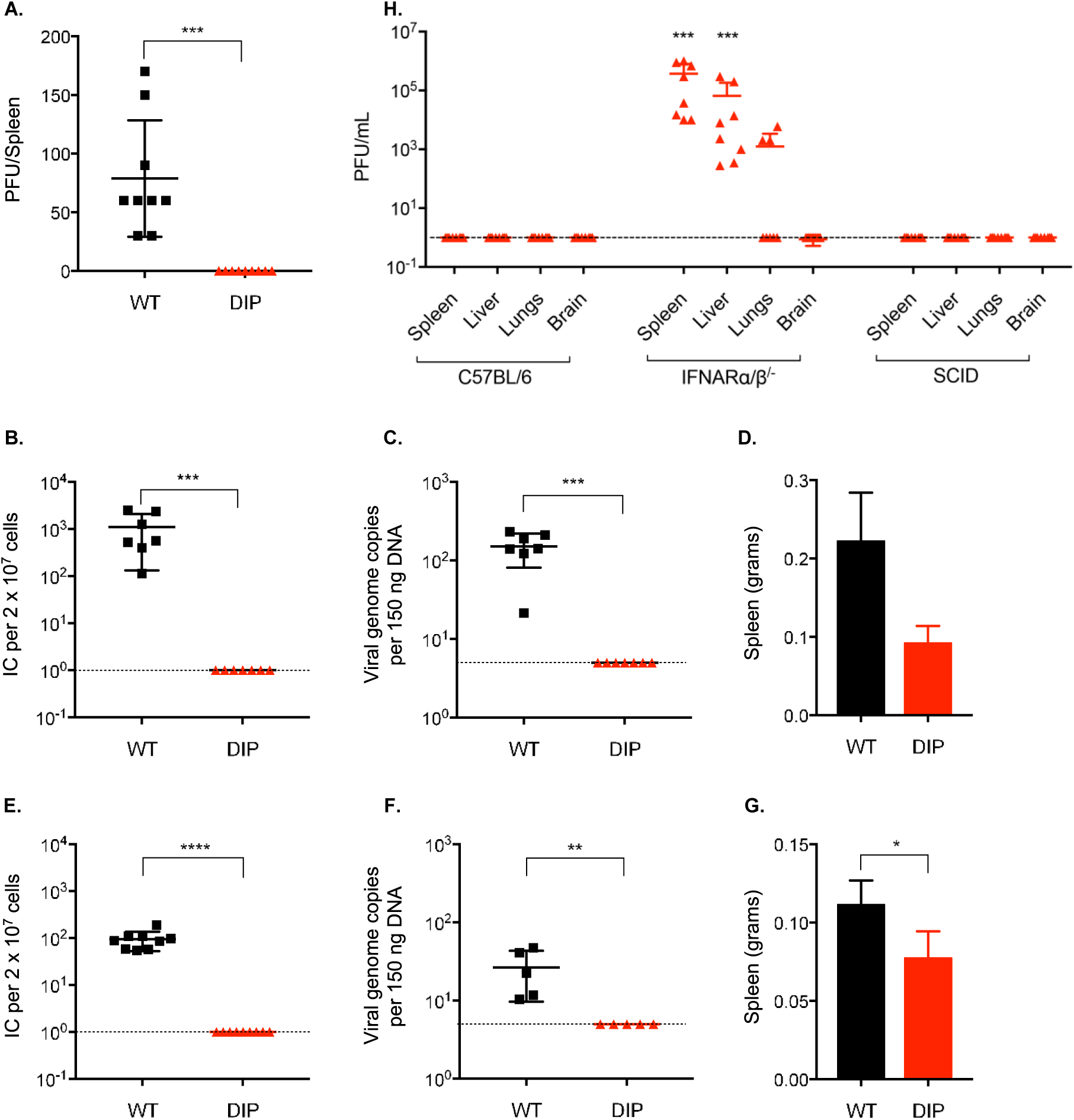
DIP produces no infectious virions and is latency deficient in vivo. All infections were performed intraperitoneally using 10^5^ PFU WT or DIP. (**A**) Productive infection in the spleens 72 h post-infection was assessed by plaque assay. (**B**) Latent infection in the spleens at 14 d post-infection was evaluated by infectious center assay and (**C**) qPCR analysis of viral DNA copy numbers. (**D**) Spleen weight at 14 d post-infection was measured. No statistically significant difference was found between WT- and DIP-infected mice. (**E**) Latent infection in the spleen at 2 mo post-infection was measured by infectious center assay and (**F**) qPCR analysis of viral DNA copy numbers. (**G**) Spleen weight at 2 mo post-infection was measured. (**H**) Spleens, livers, lungs, and brains of DIP infected C57BL/6, IFNAR^-/-^, and SCID mice were harvested at 3 d post-infection. Infectious viruses were determined by plaque assay. The graphs except (A) depicts the pooled data from 2 independent experiments using different numbers of mice for each replicate. Symbols indicate individual mice and data are means and SD. Statistical significance was determined by a two-tailed Student’s *t*-test.

Latency establishment is associated with the expansion of Vβ4-specific T-cells and splenomegaly^28–30^. At 14 d post-infection, the spleens of WT-infected mice increased to 0.22 g on average while those of DIP-infected mice weighed 0.10 g (Fig. 2D), similar to uninfected mice. At 2 mo post-infection, WT-infected mice still had significantly enlarged spleens compared to those of DIP-inoculated mice (Fig. 2G).

To determine whether the IFN-I response contributed to the attenuation of the DIP virus, we injected 10^5^ PFU of DIP intraperitoneally into the interferon-α/β receptor-deficient (IFNARα/β^-/-^) mice. DIP replication was rescued and 4 x 10^5^ infectious virions were detected in the spleens at 3d post-infection (Fig. 2H). Infectious virions were also detected in the livers and lungs but not the brains of IFNAR1^-/-^ mice. In contrast, no detectable infectious virions were recovered from severe combined immune deficiency (SCID) mice. Comprehensive analyses of the spleens, livers, brains, and lungs showed no evidence of infectious virions in either C57BL/6 or SCID mice, both of which have intact IFN-I responses (Fig. 2H).

### DIP immunization prevents latent infection

γ-herpesvirus associated malignancies are linked to latency^31,32^. Therefore, the goal of vaccination against γ-herpesviruses is to prevent latency establishment. We assessed the level of protection conferred by DIP immunization against latent infection by WT challenge. Mice were intraperitoneally injected with 1 x 10^5^ PFU DIP then intranasally challenged 1 mo later with 5,000 PFU WT. Mock immunized mice presented an average of 6 x 10^2^ infectious centers per 2 x 10^7^ splenocytes whereas no viral reactivation was detected in six of seven DIP-immunized mice 14 d after challenge (Fig. 3A). Analysis of viral copy number confirmed that DIP immunization provided protection against splenic latent infection (Fig. 3B). At 1 mo post-challenge, five of six (83.3%) DIP-vaccinated mice were completely protected from latent infection (Fig. 3C). The remaining DIP-vaccinated mouse had a 100-fold reduction in latently infected cells compared to mock immunized mice. We also challenged the immunized mice 6 mo after a single vaccination and measured viral latency at 28 d post-challenge. All six DIP-immunized mice were completely protected against latent infection by a WT virus challenge (Fig. 3D).

**Figure 3.**
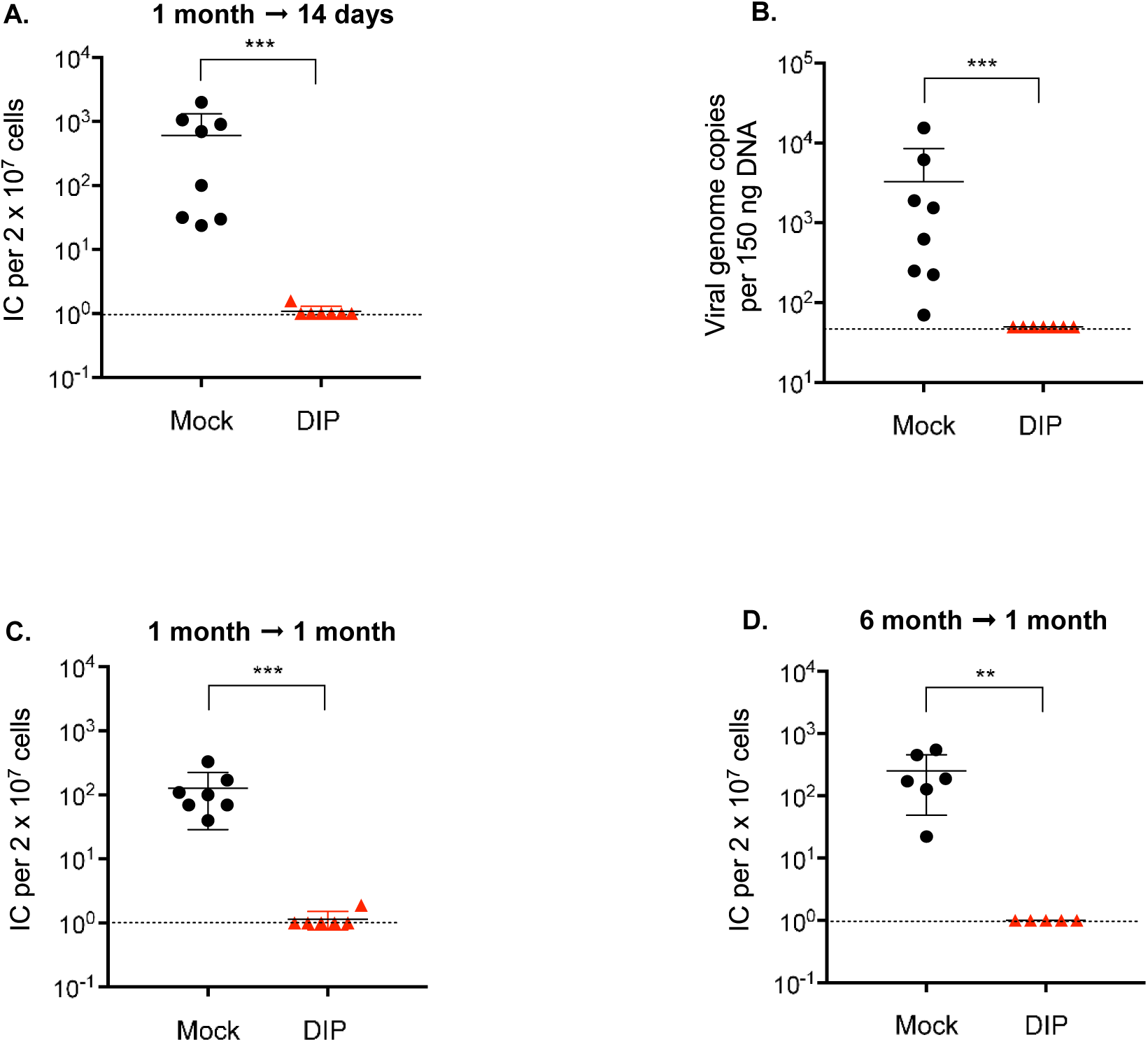
DIP vaccination confers durable protection. Mice were intraperitoneally vaccinated with 10^5^ PFU DIP and challenged intranasally with 5 x 10^3^ PFU WT virus at 1 (**A-C**) or 6 (**D**) mo post-vaccination. Latent infection in the spleen was examined at 14 (**A, B**) or 28 (**C, D**) d after challenge. Viral loads were determined by infectious center assay (**A, C, D**) and qPCR (**B**). Dotted line indicates detection limit. The graph depicts the pooled data from 2 independent experiments using different numbers of mice for each replicate. Data for individual mice, means, and SD were plotted. Statistical significance was analyzed by a two-tailed Student’s *t*-test.

### DIP primes virus-specific T cells that limit WT infection

We hypothesized that the DIP vaccine elicited a robust and functional T cell response accounting for the long-lasting protection against WT challenge. We quantified virus-specific CD8^+^ T-cells using tetramers for the MHV-68 epitopes, ORF6_487-495_ and ORF61_524-531_. At 1 mo post-infection, WT and DIP induced similar frequencies of specific T-cells to ORF6 and ORF61 (Figs. S2A & S2B). At 2 mo post-infection, the frequency of ORF6-specific T cells increased two-fold in DIP compared to WT while the frequency of ORF61-specific T-cells were similar (Figs. 4A & 4B). The effector/memory subtypes of these virus-specific T cells were examined by the expression levels of IL7Rα (CD127) and killer cell lectin-like receptor (KLRG1) (Fig. S2D). The CD127^high^KLRG1^low^ subset are memory precursors effector cells (MPECs), which develop into long-lived memory cells, whereas the CD127^low^KLRG1^high^ subset, referred to as short-lived effector T cells (SLECs), are terminally differentiated^33^. We observed that DIP promoted the generation of CD8^+^ MPECs. Significantly more ORF6-specific CD8^+^ T cells (54%) primed by DIP differentiated into MPECs compared to those primed by WT (33%). However, no difference was found between WT and DIP infection in terms of ORF61-specific T cells (Figs. 4C and S2C).

**Figure 4.**
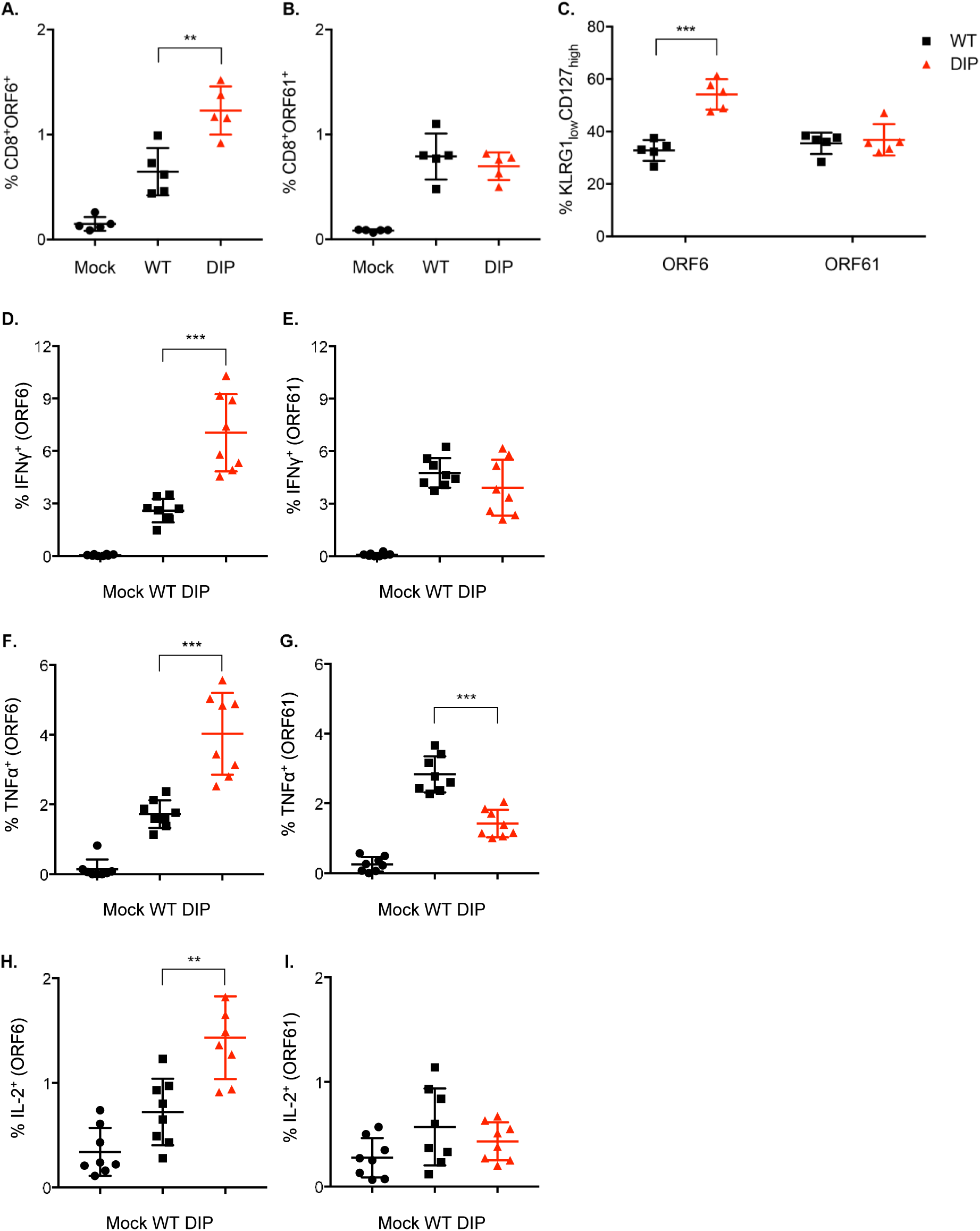
DIP elicits robust virus-specific T cell immunity. Mice were mock-infected or intraperitoneally injected with 10^5^ PFU WT or DIP. (**A, B**) At 2 mo post-infection, splenocytes were harvested and examined for virus-specific CD8+ T cells using the tetramers ORF6_487–495_/Db and ORF61_524–531_/Kb. (**C**) Tetramer-positive CD8+ T cells were examined for KLRG1 and CD127 expression. (**D-I**) Splenocytes were stimulated with ORF6_487–495_ peptide (**D, F, H**) or ORF61_524–531_ peptide (**E, G, I**) and stained for intracellular IFN-γ (**D, E**), TNF-α (**F, G**), and IL-2 (**H, I**).

We assessed the functions of these virus-specific T-cells by examining their abilities to produce IFN-γ, TNF-α, and IL-2. Consistent with the tetramer-staining results, cells producing IFN-γ, TNF-α, or IL-2 upon stimulation of the ORF6 peptide were twice as frequent in DIP-infected mice as in WT-infected mice (Figs. 4D, 4F, & 4H). Cells producing IFN-γ upon stimulation of the ORF61 peptide were at similar frequencies between WT and DIP infection. However, a lower frequency of cells primed by DIP produced TNF-α in response to the ORF61 peptide compared to those primed by WT (Figs. 4E & 4F). The ORF61 peptide did not stimulate any cells from either WT or DIP-infected mice to produce IL-2 (Fig. 4I). Therefore, despite its limited and transient antigen expression due to highly attenuated replication, DIP still induces robust and functional T-cell responses.

To determine whether DIP-primed T-cells confer protection against WT challenge, we harvested CD4^+^, CD8^+^ or total T-cells from mice infected 2 mo earlier with WT or DIP and transferred 3 x 10^6^ cells into naïve mice. These recipient mice were challenged with 5000 PFU WT 1 d after transfer. No significant difference was observed between WT- and DIP-primed T cells in terms of donor cell expansion (Fig. S3). At 14d post-challenge, CD4+ T-cell transfer had minimal impact on the number of latently infected cells (Fig. 5A) despite evidence that CD4+ T cells are cytotoxic to herpesviruses^34–36^. The transfer of WT-primed CD8+ T-cells caused a five-fold reduction in reactivated latently infected cells. In contrast, CD8+ T-cells primed by DIP failed to affect the latently infected cell pool (Fig. 5B). However, naïve mice receiving DIP-primed total T-cells had a 30-fold reduction in the number of reactivated latently infected cells, whereas transferring of WT-primed total T cells caused a 20-fold reduction (Fig. 5C). The results indicate that virus-specific CD4+ and CD8+ T-cells act cooperatively to confer protection. Despite severe attenuation, DIP vaccination elicited robust cellular immunity that inhibits the establishment of latency by the challenge virus.

**Figure 5.**
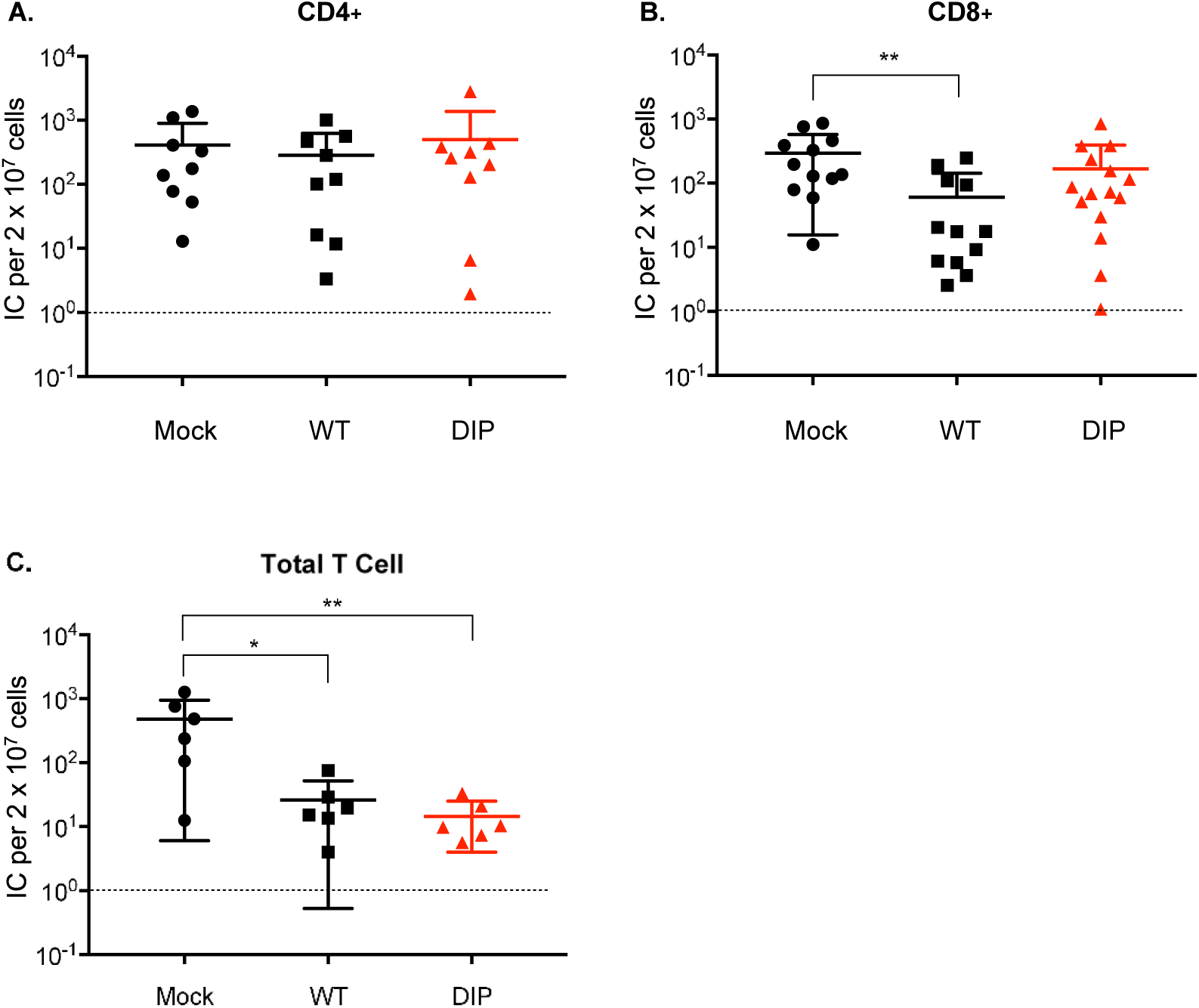
CD4+ and CD8+ T-cells confer antiviral protection. CD4+, CD8+, or total T cells were purified via negative selection from the spleens of mock-infected mice or mice that were intraperitoneally infected with 10^5^ PFU WT or DIP 2 mo previously. Three million CD4+ (**A**), CD8+ (**B**), or total T (**C**) cells were transferred to a congenic mouse by tail vein injection. The recipient mice were intranasally challenged with 5 x 10^3^ PFU WT at 24 h post-transfer. Latent infection in the spleen at 14 d post challenge was measured by infectious center assay. Pooled data from 2 independent experiments using different numbers of mice for each replicate. Data for individual mice, means, and SD were plotted. Statistical significance was analyzed by a two-tailed Student’s *t*-test.

### Optimal DIP-mediated protection requires both antibodies and T cells

To determine whether DIP-elicited antibodies complemented the T-cell-mediated protection, serum and total T-cell from DIP-infected mice were transferred to naïve mice. This combination completely protected four of six mice against a 5000 PFU WT challenge (Fig. 6A). The two unprotected mice had a significantly reduced number of reactivating latently infected cells compared to the control. We examined the protective capacity of antibodies by passively transferring DIP-immune serum to naïve mice. No significant difference in protection was observed between those receiving DIP-immune serum and those receiving serum from mock infected mice (Fig. 6B). Neutralizing activity in DIP-immune serum was less than that in WT-immune serum at 2 mo post-infection (Fig. 6D) despite relatively higher levels of virus-specific IgG in the serum DIP-infected mice (Fig. 6C). These results indicate that DIP-elicited humoral immunity collaborates with cellular immunity to provide optimal protection.

**Figure 6.**
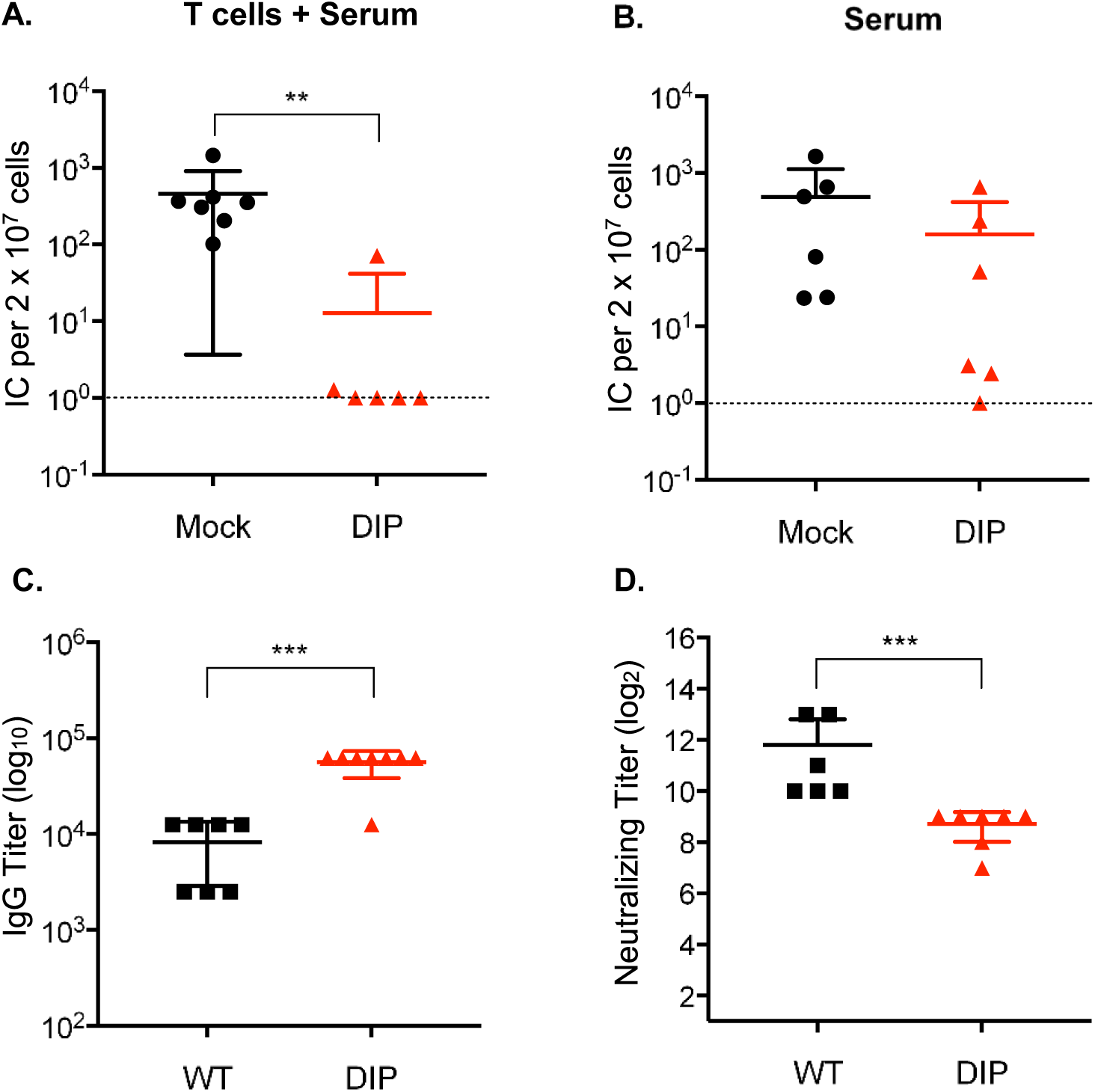
DIP vaccination elicits protective antibodies. Mice were intraperitoneally infected with 10^5^ PFU WT or DIP 2 mo previously. (**A**) Total T cells and sera isolated from mock- or DIP-infected mice were transferred to congenic naïve mice by tail vein and intraperitoneal injections, respectively. Recipient mice were intranasally challenged 24 h later with 5 x 10^3^ PFU WT virus. Latent infection in the spleen at 14 d post-challenge was assessed by infectious center assay. (**B**) Sera collected from uninfected- or DIP-infected mice were transferred to naïve mice that were intranasally challenged 24 h later with 5 x 10^3^ PFU WT virus. Latent infection in the spleen at 14 d post-challenge was evaluated by infectious center assay. (**C, D**) Sera collected from infected mice were analyzed for virus-specific IgG by ELISA and for neutralizing activity. Pooled data from 2 independent experiments using different numbers of mice for each replicate. Means and SD were plotted. Statistical significance was analyzed by a two-tailed Student’s *t*-test.

### DIP vaccine elicits robust inflammatory responses

Despite its limited replication, DIP primed robust virus-specific immune responses and conferred durable protection. Activation of innate immune response is essential for the development of adaptive immunity^54-56^. WT MHV-68 avoids inducing inflammatory cytokines in order to evade the immune system. *In vitro*, a high MOI (100 PFU/cell) was required to elicit a measurable cytokine response in bone marrow-derived macrophages (BMDM) and dendritic cells^37^. We investigated whether DIP induces inflammatory cytokines. BMDMs were infected with WT and DIP at MOI of 1 and cytokine RNA expression was quantified at 24 h post-infection. IFN-β, TNF-α, IL-6, and IL-12p40 were significantly upregulated in response to DIP infection compared to WT (Fig. 7A). In addition, WT infection at MOI of 10 still did not induce cytokine expression (Fig. S4). IL-12 is critical for Th1 polarization and cytotoxic cellular immune responses^38^. We validated the IL-12p40 RNA expression by measuring the protein with enzyme-linked immunosorbent assay (ELISA). DIP induced 30-fold more IL-12p40 protein than WT infection (Fig. 7B). The ability of DIP to stimulate the innate immune responses *in vivo* was also determined. Two days after intraperitoneal injections of viruses, there were five times as many peritoneal exudate cells (PECs) in DIP-infected as in mock-infected mice and significantly more cells than in WT-infected mice (Fig. 7C). Flow cytometry analysis of cellular compositions revealed that DIP significantly induced more plasmacytoid DCs (pDCs) than WT (Fig. 7D). As pDCs produce IFN-I^39^, we also detected the upregulation of ISG54 and IFIT2 in the PECs of DIP-infected mice compared to those of WT-infected mice (Fig. 7E). Taken together, the foregoing results indicate that DIP is highly efficacious at inducing inflammatory responses.

**Figure 7.**
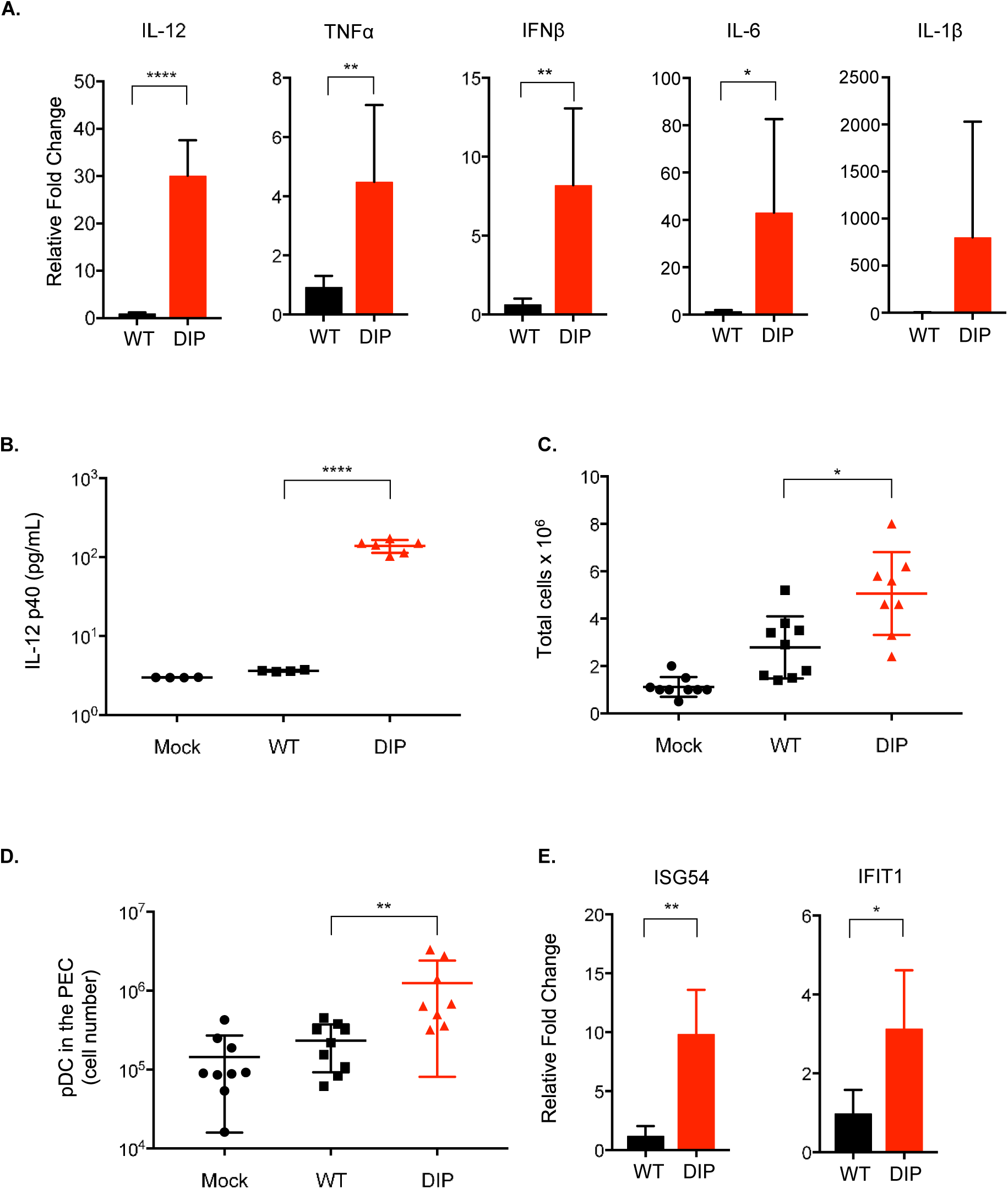
DIP elicits inflammatory and immunomodulatory cytokines. Mouse BMDMs were infected with WT or DIP at MOI = 1 (triplicate). (**A**) Total RNA was extracted 24 h post-infection for reverse transcription and qPCR to measure the expression levels of IFN-β, IL-1β, TNF-α, IL-6, IL-12, and β-actin. Cytokine RNA expression was normalized against β-actin and the relative fold change was calculated by comparison with mock-infected BMDM. (**B**) Supernatants were collected 24 h post-infection to measure IL-12p40 production by ELISA. Mice were either mock-infected or intraperitoneally injected with 10^5^ PFU WT or DIP. PECs were collected at 48 h post-infection. (**C**) Total cell numbers in the PECs were counted. (**D**) The pDCs were identified by gating on the Lin^-^(CD3^-^CD19^-^NK1.1^-^)B220^+^CD11c^Int^PDCA-1^+^ population. (**E**) Total RNA was extracted from the PECs. RNA expression of ISGs was analyzed by quantitative PCR. Means and SD were plotted. Statistical significance was analyzed by a two-tailed Student’s *t*-test.

## Discussion

An effective γ-herpesvirus vaccine should protect against the establishment of latency given the association between latent infection and tumorigenesis. Several vaccine strategies targeting single viral antigens were previously tested in the MHV-68 mouse infection model. These antigens reduced infectious mononucleosis-like symptoms of lymphoproliferation but failed to limit establishment of latency^40–45^. This finding resembles that reported for a clinical trial of EBV gp350-based vaccines^9^. The only vaccine strategy proven to reduce long-term latent viral loads in the MHV-68 model was based on live attenuated viruses designed to be latency-deficient^27,46–50^. However, a major drawback of latency-deficient viruses is the ability to undergo lytic replication^51^. In this study, we tested a strategy to attenuate the *in vivo* replication of the vaccine virus by inactivating viral antagonists of the IFN-I response. IFN-I is the first line in host antiviral defense and is critical in the development of effective immune responses. IFNs bridge innate and adaptive immunity by activating dendritic cells and inducing Th1 and potent antibody responses^52–55^. IFN-I has been tested as an adjuvant for vaccines against several distinct viruses including influenza, HIV, Ebola, CMV, and γ-herpesviruses^56–60^. Interestingly, Aricò et al. used MHV-68 to demonstrate an increase in viral-specific antibody titers when heat-inactivated virus was co-administered with IFN-α/-β^56^. We proposed that disarming viral IFN-I evasion genes may facilitate the IFN-I response, providing the adjuvanticity required for attenuated viral vaccines. In addition, we also inactivated viral inhibitor of MHC class I presentation pathway and deleted the latency locus to increase the immunogenicity and safety of the vaccine virus, DIP. The present study demonstrates that DIP is highly attenuated yet maintains overall immunogenicity relatively similar to WT.

DIP cannot undergo productive infection or persist *in vivo*. Despite its attenuated replication, DIP elicits robust innate immune responses (e.g. IL-12), memory T cells, and virus-specific antibodies with neutralizing activity. Single DIP vaccination protected against latency establishment following WT challenge. DIP-mediated protection was durable as all immunized mice remained fully protected even 6 months after a single vaccination.

Antibodies represent the first line of vaccine-mediated protection. Nevertheless, the ideal prophylactic vaccine also induces protective cellular immunity. The adoptive transfer experiments revealed that DIP-induced, virus-specific T cells and antibodies complement each other to provide optimal protection (Fig. 6A). While CD4+ T-cells alone have little protective capacity, they collaborate with CD8+ T-cells to provide protection (Fig. 5). This CD4-CD8 collaboration occurred with either WT- or DIP-primed T-cells indicating the capability of DIP to elicit effective adaptive immunity. It is recognized that CD4+ T-cells optimize the development and maintenance of memory CD8+ T-cells^61,62^. However, it is unclear whether memory CD8+ T-cells absolutely require this help from CD4+ T-cells or are simply enhanced by them^63^. The expansion of donor CD8+ T-cells in mice receiving total T cells did not surpass that in mice only receiving CD8+ T-cells (Fig. S3). Previous work indicated that memory CD4+ T-cells enhance the functionality of memory CD8+ T-cells^64–66^ but this enhancement was not examined in this study. Furthermore, the effect of memory CD4+ T-cells observed here may not have been mediated by enhancing memory CD8+ T-cells responses. Rather, CD4+ and CD8+ T-cells may target different infected cells, complementing each other, to provide effective protection against latency establishment in response to WT challenge. The mechanisms underlying the T-cell collaboration identified herein merit further investigation. It is clear that a prophylactic vaccine against γ-herpesvirus should prime both memory CD8+ and CD4+ T-cells.

A live viral vaccine induces a broad immune response against multiple viral targets especially when the mechanisms required for protection against a pathogen are not known. Furthermore, by mimicking an infection, a live vaccine stimulates multiple innate immune responses, robustly induces inflammatory and immunomodulatory cytokines, and provides adjuvanticity for long-lasting vaccine-mediated protective immunity. Nevertheless, most viruses have evolved strategies to counteract the host innate immune system. Data from BMDMs infected *in vitro* and PECs from infected mice indicates that DIP induces a stronger inflammatory response than WT (Fig. 7), partially accounting for the effective immunogenicity of DIP. DIP also recruits more pDCs than WT, which could also explain the ability of DIP to elicit robust CD8+ T-cells and antibody responses in spite of its highly attenuated replication^67–70^. Future experiments examining the role of specific cytokines or pDCs via genetic knockout and antibody depletion approaches may reveal how DIP-induced innate immune responses impact humoral and cellular immunity. Increased inflammatory cytokine production favors SLEC generation whereas shortening the duration of inflammation has been shown to accelerate MPEC development^71,72^. DIP-mediated inflammatory responses could be short-lived as DIP is highly attenuated *in vivo*. The heightened but transient DIP-induced inflammation appears to prime a robust T-cell response towards the MPEC phenotype.

The development of vaccines against human γ-herpesviruses has been hindered by their restricted host range. Neither EBV or KSHV infects small animals. While the results from mouse studies are not always directly translatable to humans, mouse models have been instrumental in elucidating fundamental principles that cannot be directly tested in humans. MHV-68 infection in mice provides a powerful, easily manipulated small animal model for analyzing fundamental events associated with the infection and immune control of γ-herpesviruses^73–78^. Moreover, the MHV-68 model serves to assess proof-of-concept vaccine strategies^79^. The results from the present study provides the guidance for a rational design of effective live EBV and KSHV vaccines that are highly attenuated and deficient in latency. Deletion of viral immune evasion genes may provide a strategy for the construction of safe yet immunogenic live vaccines against other pathogens.

## Acknowledgements

The authors thank the UCLA CFAR Virology Core (P30 AI028697) for their help with viral genome copy number analysis. We also thank the UCLA Jonsson Comprehensive Cancer Center (P30 CA016042) and the CFAR Flow Cytometry Core Facility (P30 AI028697) for assistance with flow cytometry. We thank Edward J. Usherwood of Dartmouth College and In-Jeong Kim, Kathleen G. Lanzer, and Tres Cookenham of the Trudeau Institute for scientific advice and discussion. We thank Timothy March and the UCLA Division of Laboratory Animal Medicine for veterinary assistance. We thank Autumn York and Steve J. Bensinger for providing bone marrow-derived macrophages. We would like to thank Editage (www.editage.com) for English language editing. G.B. and A.K.L were supported by an Interdisciplinary Training in Virology and Gene Therapy Training grant (No. NIH T32 AI 060567). The study was supported by NIDCR DE023591 (R01) and NCI CA177322 (P01).

## Author contributions

G.B., N.F., R.S., and T.T.W. conceived and planned the experiments. G.B. and N.F. carried out the experiment with the help of A.S., A.K.L., W.W.L., L.T., Y.H.K., and T.H. G.B., N.F., R.S., and T.T.W. analyzed and interpreted the results. C.F.W., and M.A.B. provided critical scientific advice to the research described in this manuscript. G.B., N.F., and T.T.W. wrote the manuscript with inputs from A.K.L., W.W.L., M.A.B., C.F.W., and R.S.

## Materials & Methods

### Viruses and cells

WT MHV-68 was obtained from the American Type Culture Collection (ATCC; Vr1465; Manassas, VA, USA). WT and DIP viruses were propagated in 3T3 and Vero cells and titered by plaque assay. Viruses were concentrated by high-speed centrifugation and resuspended in serum-free Dulbecco’s modified Eagle’s medium (DMEM). Vero cells were cultured in DMEM containing 10% (w/v) fetal bovine serum (FBS) supplemented with penicillin and streptomycin. The 3T3 cells were cultured in DMEM containing 10% (w/v) bovine calf serum (BCS) and 1% penicillin and streptomycin.

### Plaque assay

Each sample was serially diluted tenfold and incubated on Vero cells on 12-well plates in duplicate. The inoculum was removed after 1 h of incubation and the cells were overlaid with 1% (w/v) methylcellulose in DMEM containing 10% (w/v) FBS. Six days post-infection, the cells were fixed with 2% (w/v) crystal violet in 20% (v/v) ethanol. Viral titers were determined by counting plaque numbers. To determine viral titers in the mouse tissues, 1-mL homogenates were prepared in a Dounce homogenizer (Thomas Scientific, Swedesboro, NJ, USA) and used for the plaque assay. Plaques were counted and viral titers in each tissue were expressed in PFU mL^-1^.

### In vitro growth curve

The 3T3 cells were plated on media with or without IFN-β (100 U mL^-1^) for 24 h. Cells were infected at MOI = 0.01 with WT or DIP virus for 1 h at 37 °C. The inoculum was then removed and the cells were washed twice with media before adding fresh media with or without IFN-β (100 U mL^-1^). Cells and supernatant were harvested 24 h, 48 h, and 72 h post-infection for the plaque assay.

### Construction of DIP vaccine

The recA+ *Escherichia coli* GS500 harboring a BAC containing the WT MHV-68 genome was used to construct recombinant MHV-68 by allelic exchange with conjugation-competent *E. coli* GS111 containing the suicide shuttle plasmid pGS284^74-76^. For each recombinant MHV-68, an overlap extension PCR was used to construct the unique shuttle plasmid pGS284 harboring the desired mutation and a ∼500-bp flanking region. Sequences upstream of the desired mutation (A fragments) were amplified by AF and AR primers. The downstream sequences (B fragments) were amplified by BF and BR primers using wild type MHV-68 virion DNA as the template. The A and B fragments had > 20-bp overlapping sequences. For the subsequent PCR reaction, the A and B fragments were used as templates and amplified by AF and BR primers. The final PCR products were digested with the appropriate enzymes and cloned into pGS284. To screen for the correct mutation, restriction enzyme digestion was performed on the PCR products obtained using the AF and BR primers on the BAC MHV-68 clones. Sequential allelic exchanges were conducted to obtain the final recombinant clone containing all the designed mutations (Fig. 1). After the desired recombinant clone was selected, the MHV-68 BAC was purified and transiently transfected with Lipofectamine™2000 into 293T cells with equal amounts of plasmid expressing Cre recombinase to remove the BAC sequence. Three days post transfection, a single viral clone was isolated by limiting dilution. It was then propagated for use in subsequent experiments. The viruses were quantified by plaque assay and limiting dilution.

The ORF36 and ORF54 primers were used to construct the shuttle plasmids^18,80^. Primers used to construct the other shuttle plasmids are listed in Supplementary Table 1. Primers 1-8 were used to construct shuttle plasmids for the stop codon mutation. To construct the shuttle plasmid to replace the latency locus with RTA expression driven by the PGK promoter, four fragments were amplified with primers 9-16, A (ORF72), B (RTA coding sequence and poly A tail), C (PGK promoter), and D (ORF74). The ABCD fused fragment was then generated to be cloned into pGS284.

### Mice

The animal studies were approved by the Animal Research Committee at the University of California, Los Angeles (UCLA), Los Angeles, CA, USA. Female C57BL/6J, SCID, and B6.SJL-*Ptprc*^*a*^ *Pepc*^*b*^/BoyJ mice were obtained from Jackson Laboratory, Bar Harbor, ME, USA. IFNAR^-/-^ mice were donated by Genhong Cheng at UCLA. Mice aged 6-8 wks were intraperitoneally infected with 10^5^ PFU virus in 200 μL. Intranasal vaccinations and challenges were performed by anesthetizing the mice with isoflurane and administering 20 μL virus dropwise. At the endpoint, mice were euthanized and their tissues were collected in 1 mL DMEM and homogenized with mesh filters and a Dounce homogenizer. Tissue lysates were clarified by centrifugation and used in the plaque assays. Their DNA was extracted with a DNeasy blood and tissue kit (Cat. No. 69504; Qiagen, Hilden, Germany). For the infectious center assay and the flow cytometry study, single-cell suspensions were obtained from the spleens and the red blood corpuscles were lysed in ACK (ammonium-chloride-potassium) buffer.

### Phenotyping virus-specific T cells

Before staining, the splenocytes were incubated with FC block (No. 553142; BD Bioscience, Franklin Lakes, NJ, USA). Tetramers were obtained from the NIH Tetramer Core Facility, Atlanta, GA, USA. Allophycocyanin-conjugated MHCI tetramers specific for the MHV68 epitopes D^b^/ORF6487–495 (AGPHNDMEI), K^b^/ ORF61524–531 (TSINFVKI), K^b^/ORF75c940–947 (KSLTYYKL), and K^b^/ ORF8_604-612_ (KNYIFEEKL) were incubated with splenocytes for 1 h at room temperature. Surface-staining with the following antibodies was performed by incubation at 4 °C for 30 min: anti-KLRG1 (No. 46-5893; eBioscience/Affymetrix, Santa Clara, CA, USA), anti-CD127 (No. 17-1273; eBioscience/Affymetrix, Santa Clara, CA, USA), anti-CD8 (No. 48-0081; eBioscience/Affymetrix, Santa Clara, CA, USA), anti-CD4 (No. 11-0042; eBioscience/Affymetrix, Santa Clara, CA, USA), anti-CD3 (No. 25-0031; eBioscience/Affymetrix, Santa Clara, CA, USA), anti-CD44 (No. 11-0441; eBioscience/Affymetrix, Santa Clara, CA, USA), anti-CD62L (No. 83-062; eBioscience/Affymetrix, Santa Clara, CA, USA), anti-CCR7 (No. 47-1971; eBioscience/Affymetrix, Santa Clara, CA, USA), anti-CD45.1 (No. 47-0453; eBioscience/Affymetrix, Santa Clara, CA, USA), and anti-CD45.2 (No. 12-0454; eBioscience/Affymetrix, Santa Clara, CA, USA). For intracellular staining, BD Cytofix and Cytoperm (Cat. No. 554714; BD Bioscience, Franklin Lakes, NJ, USA) were used before incubating splenocytes with anti-IFN-γ (No. 17-7311; eBioscience/Affymetrix, Santa Clara, CA, USA), anti-TNF-α (No. 46-7321; eBioscience/Affymetrix, Santa Clara, CA, USA), and anti-IL-2 (No. 25-7021; eBioscience/Affymetrix, Santa Clara, CA, USA) antibodies at room temperature for 30 min. All samples were fixed in 1% (w/v) paraformaldehyde (PFA). All experiments were analyzed on a SORP BD LSRII analytic flow cytometer (BD Bioscience, Franklin Lakes, NJ, USA). Data were analyzed in FlowJo (FlowJo LLC, Ashland, OR, USA).

### Ex vivo T cell peptide stimulation

B cells in splenocytes were depleted by incubation in flasks coated with AffiniPure goat anti-mouse IgG (H+L) (Jackson ImmunoResearch Laboratories Inc., West Grove, PA, USA) for 1 h at 37 °C. B-cell-depleted splenocytes from infected mice (CD45.2+) were incubated with naïve splenocytes (CD45.1+) at a 1:1 ratio in culture media containing 10 U mL^-1^ IL-12, 10 μg mL^-1^ brefeldin A, and 1 μg mL^-1^ peptide for 5 h at 37 °C. Splenocytes were stained and processed for flow cytometry with the indicated tetramers and surface marker antibodies.

### Infectious center assay

Serially diluted splenocytes were plated on a Vero cell monolayer and incubated overnight at 37 °C. The splenocytes were aspirated then washed off by gentle agitation. The Vero cells were overlaid with 1% (w/v) methylcellulose in DMEM containing 10% (w/v) FBS for 6 d before fixing with 2% (w/v) crystal violet in 20% (v/v) ethanol. Infectious centers indicated by plaques were counted.

### Quantitative PCR (qPCR)

The qPCR was performed on MJ Opticon 2 using PerfeCTA Fastmix (Quantabio, Beverly, MA, USA). For the viral genome copy number analysis, 150 ng extracted DNA (∼2 x 10^4^ cells) and the primers were annealed to the upstream of the ORF6 coding sequence (ORF6: 5’-TGCAGACTCTGAAGTGCTGACT-3’ and 5’-ACGCGACTAGCATGAGGAGAAT-3’) were used.

For the RNA expression analysis, cells were harvested in TRIzol (Thermo Fisher Scientific, Waltham, MA, USA) for RNA extraction according to the recommended protocol. Total RNA was treated with DNAse and used for reverse transcription in a qScript cDNA synthesis kit (Quantabio, Beverly, MA, USA) to generate cDNA for qPCR.

### Gene expression analysis by qPCR

Cell lysates were stored in TRIzol at −80 °C. Isolated RNA was treated with DNase I then used to generate cDNA in a qScript cDNA synthesis kit (Quantabio, Beverly, MA, USA) followed by gene expression analysis with PerfeCTa Fastmix (Quantabio, Beverly, MA, USA). The primers used in qPCR for IL-1β, TNF-α, IL-6, IL-12, and β-actin are listed in Supplementary Table 2.

### Infection of BMDM

Cells were harvested from bone marrow and differentiated into macrophages (BMDM) by incubation for 7 d in DMEM containing 20% (w/v) FBS, 5% (w/v) M-CSF, 1% (w/v) penicillin and streptomycin, 1% (w/v) glutamine, and 0.5% (w/v) sodium pyruvate. The BMDMs were infected with WT or DIP at MOI = 1. At 24 h post-infection, total RNA was extracted with TRIzol. Supernatants were collected for analysis in an IL-12/IL-23 p40 (total) mouse uncoated ELISA kit (No. 88-7120-22; Thermo Fisher Scientific, Waltham, MA, USA).

### Neutralizing activity

Twofold serially-diluted serum was incubated with 100 PFU WT virus for 1 h at 37 °C. The mixture was plated on a Vero cell monolayer for 1 h at 37 °C then removed. The plate was overlaid with 1% (w/v) methylcellulose in DMEM containing 10% (w/v) FBS for 6 d before fixing with 2% (w/v) crystal violet in 20% (v/v) ethanol. The neutralizing titer was taken as the highest dilution maintaining the ability of the diluted serum to reduce the number of plaques by 50% relative to the virus mixture containing fourfold diluted mock serum.

### Virus-specific IgG ELISA

A 5 μg mL^-1^ WT virion antigen solution coated a 96-well plate which was then incubated overnight at 4 °C. The plate was blocked overnight in PBS containing 1% (w/v) BSA and 0.05% (w/v) Tween-20. The plate was washed twice with PBS-T (PBS containing 0.5% Tween-20). Mouse sera were diluted in ELISA buffer (PBS containing 0.1% BSA and 0.025% Tween-20) and incubated on the plate for 1 h at room temperature. The plate was washed thrice with PBS-T and then once with PBS. A substrate solution consisting of one tablet each of *o*-phenylenediamine and urea hydrogen peroxide (No. P9187; Sigma-Aldrich Corp., St. Louis, MO, USA) in 10 mL ddH_2_O, was added to the plate. The plate was incubated for 30 min in the dark at 4 °C. The reaction was stopped by adding 4N H_2_SO_4_ and the plate was read at 490 nm and 620 nm. The virus-specific IgG titer was taken as the highest dilution generating signals higher than those of the 1:50 diluted mock serum.

### Serum transfer

Sera were obtained from mice at 2 mo post-infection. Then 200 μL pooled heat-inactivated serum was intraperitoneally injected into naïve mice. After 24 h, the naïve recipient mice were challenged intranasally with 5 x 10^3^ PFU WT. A second dose of 200 μL pooled heat-inactivated serum was intraperitoneally injected 7 d after the WT challenge. Splenocytes were harvested 14 d after the challenge for the infectious center assay.

### Adoptive T cell transfer

Splenocytes were isolated from mice at 2 mo post-infection and pooled from multiple mice. Splenocytes were negatively selected for CD4, CD8, or total T cells using EasySep isolation kits (Catalog Nos. 19765, 19853, and 19851; STEMCELL Technologies Inc., Vancouver, BC, Canada). Negative selection was confirmed by flow cytometry analysis to > 90% purity. Three million cells in 100 μL were injected into the tail vein of each B6.SJL-*Ptprc*^*a*^ *Pepc*^*b*^/BoyJ mouse. Twenty-four hours after T-cell transfer, the recipient mice were intranasally challenged with 5 x 10^3^ PFU WT MHV-68. Spleens were harvested 14 d post-challenge for the infectious center assay and flow cytometry to confirm the presence of donor T-cells in the recipient mice with anti-CD45.1 and anti-CD45.2.

### Statistical analysis

Data are presented as means and their differences were analyzed by a two-tailed unpaired Student’s *t*-test unless otherwise indicated. *P* < 0.05*, *P* < 0.01**, *P* < 0.001***, and *P* < 0.0001****.

## Supplementary Data

**Figure S1.**
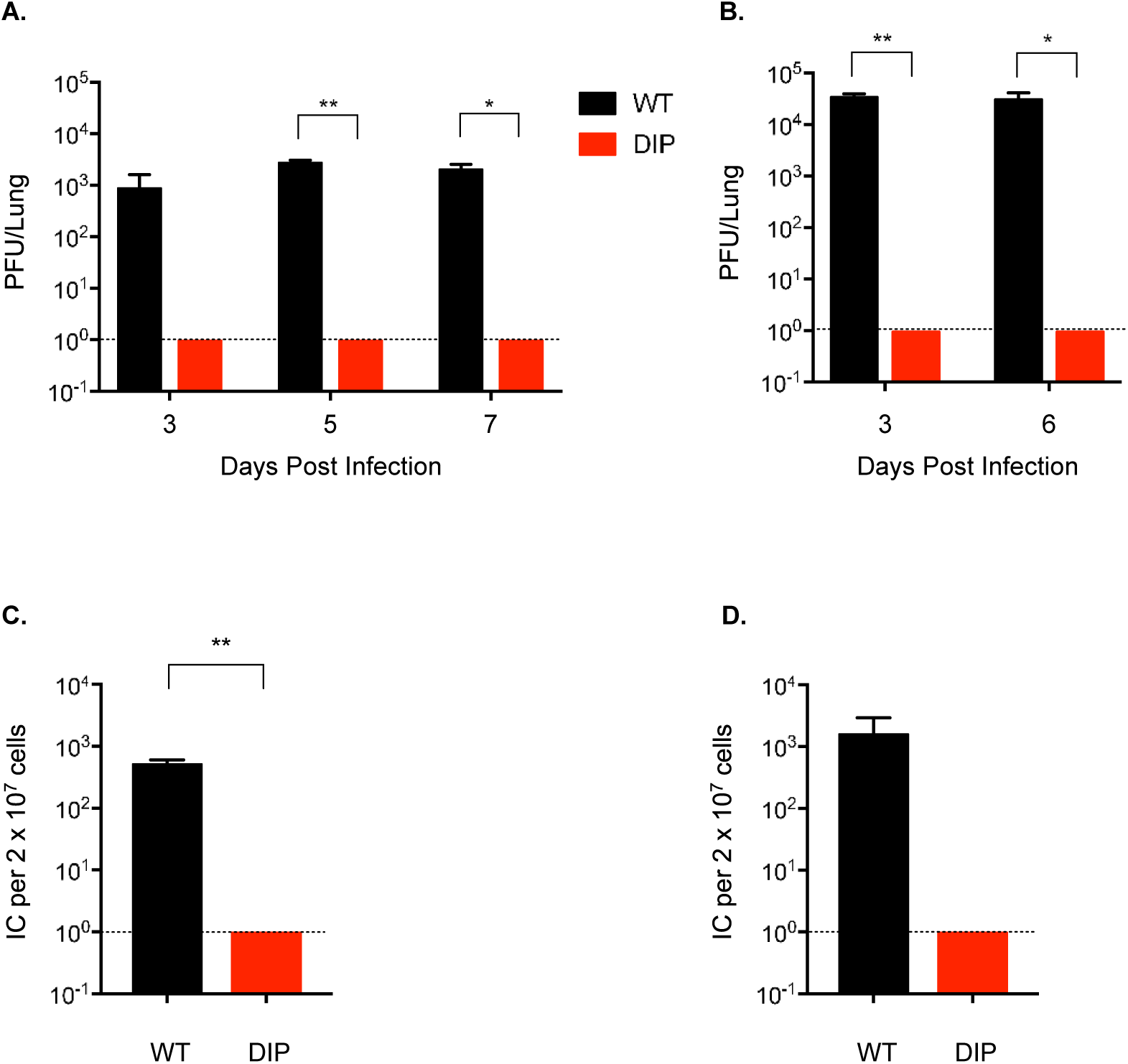
In vivo DIP virus is both replication- and latency deficient upon intranasal inoculation. Mice were intranasally inoculated with 5,000 (**A, C**) or 10^5^ PFU (**B, D**) WT or DIP. (**A, B**) Lungs (n = 3) were excised at the times indicated at the bottoms of the graphs for plaque assay. (**C, D**) Spleens (n = 3) were excised 14 d post infection for infectious center assay. Means and SD were plotted. Statistical significance was analyzed by a two-tailed Student’s *t*-test.

**Figure S2.**
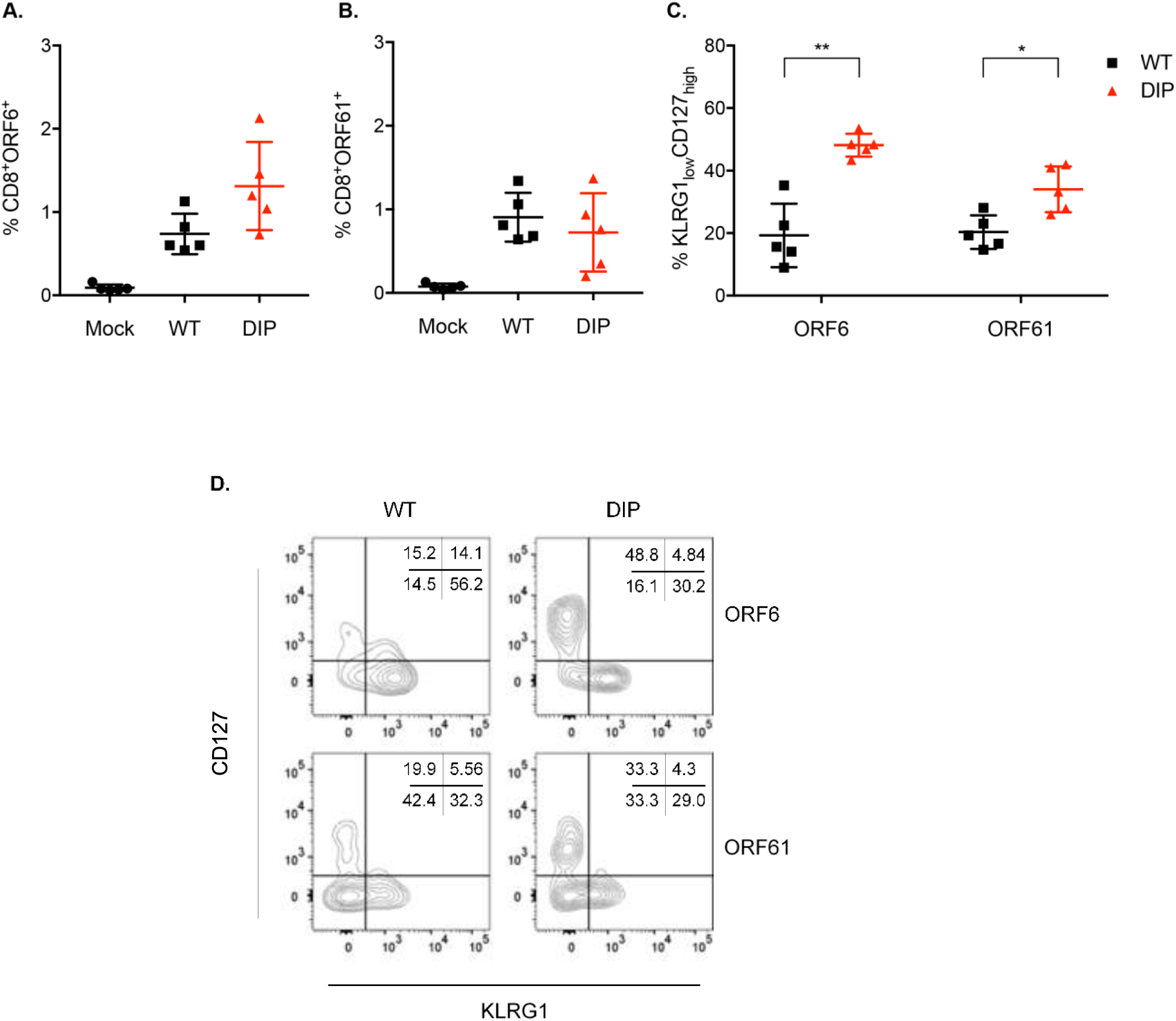
DIP infection induces a robust, virus-specific T cell response. Mice were either mock-infected or intraperitoneally infected with 10^5^ PFU WT or DIP. (**A, B**) At 1 mo post-infection, splenocytes were harvested and examined for virus-specific CD8+ T cells using the tetramers ORF6_487–495_/Db and ORF61_524–531_/Kb. (**C**) Tetramer-positive CD8^+^ T cells were examined for KLRG1 and CD127 expression. Data for individual mice (n = 5), means, and SD were plotted. Statistical significance was analyzed by a two-tailed Student’s *t*-test. (**D**) Gating strategy to determine MPEC and SLEC population frequencies using representative samples from WT- and DIP-inoculated mice.

**Figure S3.**
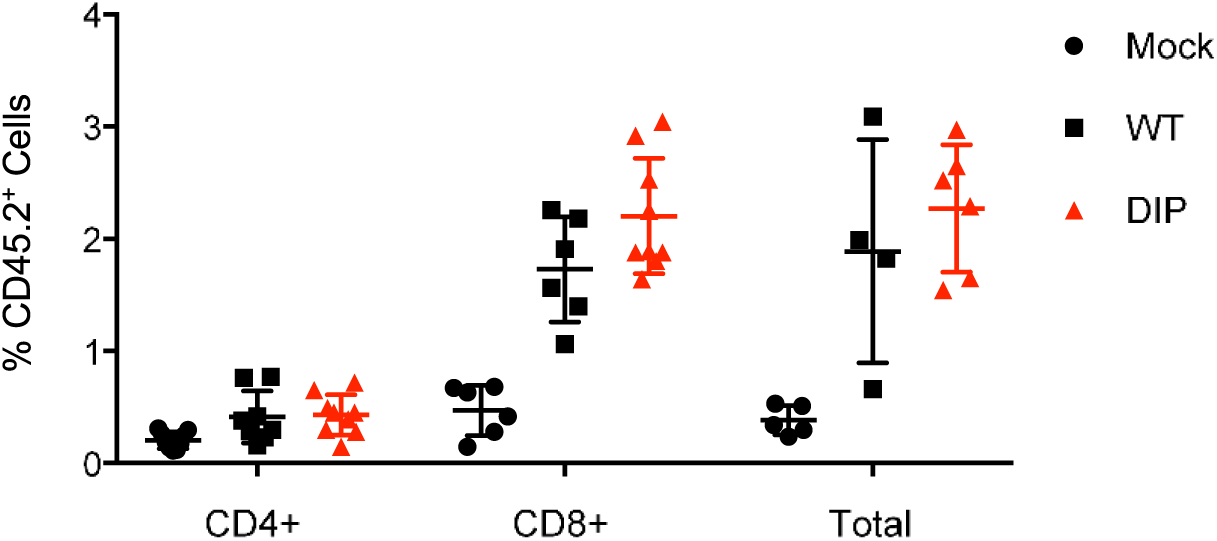
WT- and DIP-primed T cells expand to the same extent in the recipient mice. CD4^+^, CD8^+^, or total T cells were purified via negative selection from the spleens of mock-infected mice or mice intraperitoneally infected with 10^5^ PFU WT or DIP. Three million purified cells were transferred to a congenic mouse by tail vein injection. At 14 d after transfer, donor cells were analyzed by flow cytometry and the percentages are shown. Data for individual mice, means, and standard deviations are plotted. Statistical significance was analyzed by a two-tailed Student’s *t*-test. No statistical significance was determined for the percentages of WT- and DIP-primed donor cells.

**Figure S4.**
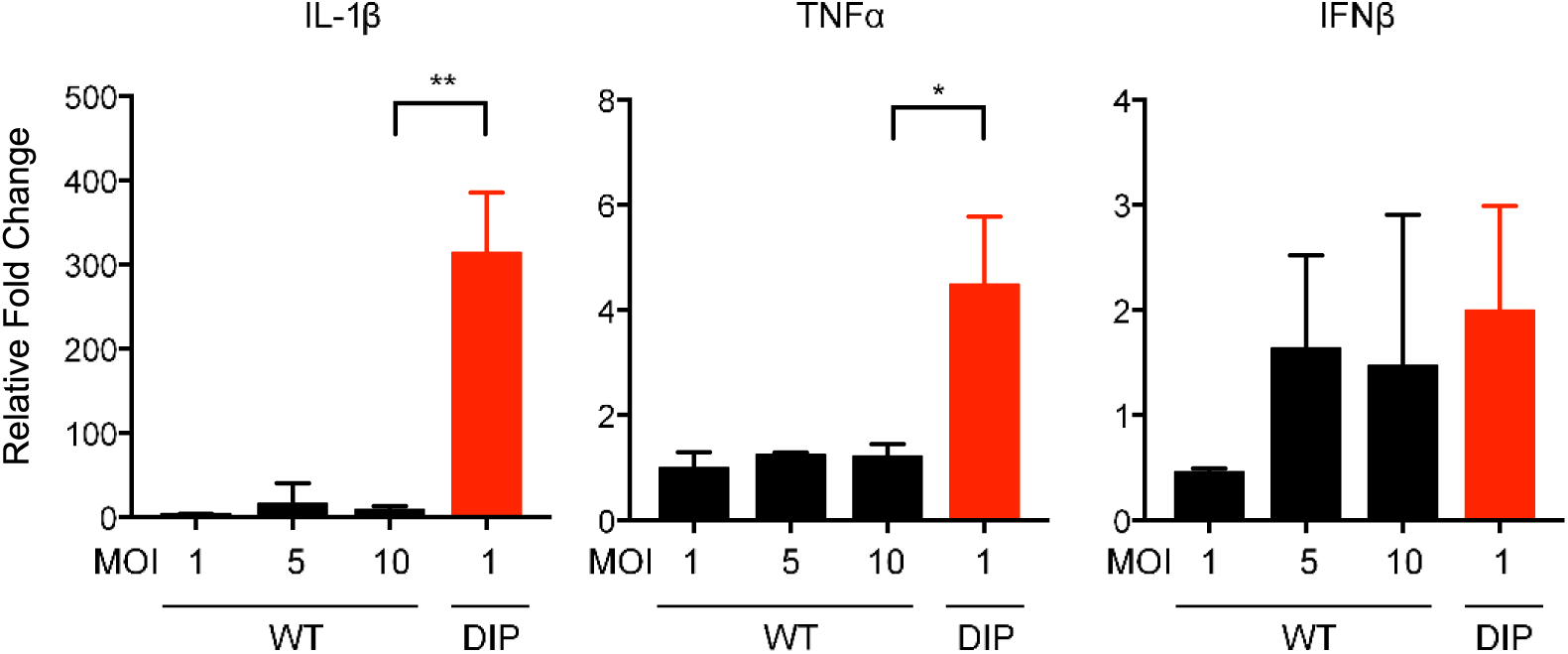
WT does not upregulate inflammatory cytokines to the same level as DIP. Mouse BMDM were infected with WT or DIP at the MOIs indicated at the bottoms of the graphs (triplicate). (**A**) Total RNAs were extracted 24 h post-infection to measure IFNβ, IL-1β, TNF-α, and β-actin expression. Cytokine RNA expression was normalized against β-actin. Relative fold change was calculated by comparing to mock-infected BMDM. Statistical significance was analyzed by a two-tailed Student’s *t*-test.

## References

1. Epstein, M. A., Achong, B. G. & Barr, Y. M. VIRUS PARTICLES IN CULTURED LYMPHOBLASTS FROM BURKITT’S LYMPHOMA. Lancet 1, 702–703 (1964).

2. zur Hausen, H. et al. EBV DNA in biopsies of Burkitt tumours and anaplastic carcinomas of the nasopharynx. Nature 228, 1056–1058 (1970).

3. Kutok, J. L. & Wang, F. Spectrum of Epstein-Barr virus-associated diseases. Annu Rev Pathol 1, 375–404 (2006).

4. Chang, Y. et al. Identification of herpesvirus-like DNA sequences in AIDS-associated Kaposi’s sarcoma. Science 266, 1865–1869 (1994).

5. Cesarman, E., Chang, Y., Moore, P. S., Said, J. W. & Knowles, D. M. Kaposi’s sarcoma-associated herpesvirus-like DNA sequences in AIDS-related body-cavity-based lymphomas. N. Engl. J. Med. 332, 1186–1191 (1995).

6. Soulier, J. et al. Kaposi’s sarcoma-associated herpesvirus-like DNA sequences in multicentric Castleman’s disease. Blood 86, 1276–1280 (1995).

7. Plummer, M. et al. Global burden of cancers attributable to infections in 2012: a synthetic analysis. The Lancet Global Health 4, e609–e616 (2016).

8. Thorley-Lawson, D. A. & Poodry, C. A. Identification and isolation of the main component (gp350-gp220) of Epstein-Barr virus responsible for generating neutralizing antibodies in vivo. J. Virol. 43, 730–736 (1982).

9. Sokal, E. M. et al. Recombinant gp350 vaccine for infectious mononucleosis: a phase 2, randomized, double-blind, placebo-controlled trial to evaluate the safety, immunogenicity, and efficacy of an Epstein-Barr virus vaccine in healthy young adults. J. Infect. Dis. 196, 1749–1753 (2007).

10. Johnston, C., Gottlieb, S. L. & Wald, A. Status of vaccine research and development of vaccines for herpes simplex virus. Vaccine 34, 2948–2952 (2016).

11. Ivashkiv, L. B. & Donlin, L. T. Regulation of type I interferon responses. Nature Reviews Immunology 14, 36–49 (2014).

12. McNab, F., Mayer-Barber, K., Sher, A., Wack, A. & O’Garra, A. Type I interferons in infectious disease. Nat. Rev. Immunol. 15, 87–103 (2015).

13. Stark, G. R., Kerr, I. M., Williams, B. R. G., Silverman, R. H. & Schreiber, R. D. HOW CELLS RESPOND TO INTERFERONS. Annual Review of Biochemistry 67, 227–264 (1998).

14. Wilson, E. B. & Brooks, D. G. Decoding the complexity of type I interferon to treat persistent viral infections. Trends Microbiol. 21, 634–640 (2013).

15. Lee, H.-R., Lee, S., Chaudhary, P. M., Gill, P. & Jung, J. U. Immune evasion by Kaposi’s sarcoma-associated herpesvirus. Future Microbiol 5, 1349–1365 (2010).

16. Coscoy, L. Immune evasion by Kaposi’s sarcoma-associated herpesvirus. Nat. Rev. Immunol. 7, 391–401 (2007).

17. Virgin, H. W. et al. Complete sequence and genomic analysis of murine gammaherpesvirus 68. J. Virol. 71, 5894–5904 (1997).

18. Leang, R. S. et al. The Anti-interferon Activity of Conserved Viral dUTPase ORF54 is Essential for an Effective MHV-68 Infection. PLoS Pathogens 7, e1002292 (2011).

19. Ishido, S., Wang, C., Lee, B. S., Cohen, G. B. & Jung, J. U. Downregulation of major histocompatibility complex class I molecules by Kaposi’s sarcoma-associated herpesvirus K3 and K5 proteins. J. Virol. 74, 5300–5309 (2000).

20. Stevenson, P. G. et al. K3-mediated evasion of CD8(+) T cells aids amplification of a latent gamma-herpesvirus. Nat. Immunol. 3, 733–740 (2002).

21. Ballestas, M. E., Chatis, P. A. & Kaye, K. M. Efficient persistence of extrachromosomal KSHV DNA mediated by latency-associated nuclear antigen. Science 284, 641–644 (1999).

22. Fowler, P., Marques, S., Simas, J. P. & Efstathiou, S. ORF73 of murine herpesvirus-68 is critical for the establishment and maintenance of latency. J. Gen. Virol. 84, 3405–3416 (2003).

23. Moorman, N. J., Willer, D. O. & Speck, S. H. The Gammaherpesvirus 68 Latency-Associated Nuclear Antigen Homolog Is Critical for the Establishment of Splenic Latency. Journal of Virology 77, 10295–10303 (2003).

24. Sun, R. et al. A viral gene that activates lytic cycle expression of Kaposi’s sarcoma-associated herpesvirus. Proc Natl Acad Sci USA 95, 10866 (1998).

25. Wu, T. T., Usherwood, E. J., Stewart, J. P., Nash, A. A. & Sun, R. Rta of murine gammaherpesvirus 68 reactivates the complete lytic cycle from latency. J. Virol. 74, 3659–3667 (2000).

26. Wu, T. T., Tong, L., Rickabaugh, T., Speck, S. & Sun, R. Function of Rta is essential for lytic replication of murine gammaherpesvirus 68. J. Virol. 75, 9262–9273 (2001).

27. Jia, Q. et al. Induction of protective immunity against murine gammaherpesvirus 68 infection in the absence of viral latency. J. Virol. 84, 2453–2465 (2010).

28. Coppola, M. A. et al. Apparent MHC-independent stimulation of CD8+ T cells in vivo during latent murine gammaherpesvirus infection. J. Immunol. 163, 1481–1489 (1999).

29. Tripp, R. A. et al. Pathogenesis of an infectious mononucleosis-like disease induced by a murine gamma-herpesvirus: role for a viral superantigen? J. Exp. Med. 185, 1641–1650 (1997).

30. Usherwood, E. J., Ross, A. J., Allen, D. J. & Nash, A. A. Murine gammaherpesvirus-induced splenomegaly: a critical role for CD4 T cells. J. Gen. Virol. 77 (Pt 4), 627–630 (1996).

31. Damania, B. & Cesarman, E. Kaposi’s Sarcoma-Associated Herpesvirus. in Fields Virology (eds. Fields, B., Knipe, D. & Howley, P.) 2, 2080–2128 (Wolters Kluwer Health/Lippincott Williams & Wilkins, 2013).

32. Longnecker, R., Kieff, E. & Cohen, J. Epstein-Barr virus. in Fields Virology (eds. Fields, B., Knipe, D. & Howley, P.) 2, 1898–1959 (Wolters Kluwer Health/Lippincott Williams & Wilkins, 2013).

33. Joshi, N. S. et al. Inflammation Directs Memory Precursor and Short-Lived Effector CD8+ T Cell Fates via the Graded Expression of T-bet Transcription Factor. Immunity 27, 281–295 (2007).

34. Freeman, M. L. et al. CD4 T Cells Specific for a Latency-Associated γ-Herpesvirus Epitope Are Polyfunctional and Cytotoxic. The Journal of Immunology 193, 5827–5834 (2014).

35. Long, H. M. et al. CD4+ T-cell responses to Epstein-Barr virus (EBV) latent-cycle antigens and the recognition of EBV-transformed lymphoblastoid cell lines. J. Virol. 79, 4896–4907 (2005).

36. Stuller, K. A. & Flano, E. CD4 T Cells Mediate Killing during Persistent Gammaherpesvirus 68 Infection. Journal of Virology 83, 4700–4703 (2009).

37. Sun, C. et al. Evasion of innate cytosolic DNA sensing by a gammaherpesvirus facilitates establishment of latent infection. J. Immunol. 194, 1819–1831 (2015).

38. Trinchieri, G. Interleukin-12 and the regulation of innate resistance and adaptive immunity. Nat. Rev. Immunol. 3, 133–146 (2003).

39. Siegal, F. P. et al. The nature of the principal type 1 interferon-producing cells in human blood. Science 284, 1835–1837 (1999).

40. Obar, J. J. et al. T-Cell Responses to the M3 Immune Evasion Protein of Murid Gammaherpesvirus 68 Are Partially Protective and Induced with Lytic Antigen Kinetics. Journal of Virology 78, 10829–10832 (2004).

41. Woodland, D. L. et al. Vaccination against murine gamma-herpesvirus infection. Viral Immunol. 14, 217–226 (2001).

42. Usherwood, E. J., Ward, K. A., Blackman, M. A., Stewart, J. P. & Woodland, D. L. Latent antigen vaccination in a model gammaherpesvirus infection. J. Virol. 75, 8283–8288 (2001).

43. Stewart, J. P., Micali, N., Usherwood, E. J., Bonina, L. & Nash, A. A. Murine gammaherpesvirus 68 glycoprotein 150 protects against virus-induced mononucleosis: a model system for gamma-herpesvirus vaccination. Vaccine 17, 152–157 (1999).

44. Stevenson, P. G., Belz, G. T., Castrucci, M. R., Altman, J. D. & Doherty, P. C. A -herpesvirus sneaks through a CD8+ T cell response primed to a lytic-phase epitope. Proceedings of the National Academy of Sciences 96, 9281–9286 (1999).

45. Liu, L., Usherwood, E. J., Blackman, M. A. & Woodland, D. L. T-cell vaccination alters the course of murine herpesvirus 68 infection and the establishment of viral latency in mice. J. Virol. 73, 9849–9857 (1999).

46. Tibbetts, S. A., McClellan, J. S., Gangappa, S., Speck, S. H. & Virgin, H. W. Effective vaccination against long-term gammaherpesvirus latency. J. Virol. 77, 2522–2529 (2003).

47. Fowler, P. & Efstathiou, S. Vaccine potential of a murine gammaherpesvirus-68 mutant deficient for ORF73. J. Gen. Virol. 85, 609–613 (2004).

48. Rickabaugh, T. M. et al. Generation of a latency-deficient gammaherpesvirus that is protective against secondary infection. J. Virol. 78, 9215–9223 (2004).

49. Boname, J. M., Coleman, H. M., May, J. S. & Stevenson, P. G. Protection against wild-type murine gammaherpesvirus-68 latency by a latency-deficient mutant. J. Gen. Virol. 85, 131–135 (2004).

50. May, J. S., Coleman, H. M., Smillie, B., Efstathiou, S. & Stevenson, P. G. Forced lytic replication impairs host colonization by a latency-deficient mutant of murine gammaherpesvirus-68. J. Gen. Virol. 85, 137–146 (2004).

51. Freeman, M. L. et al. Importance of antibody in virus infection and vaccine-mediated protection by a latency-deficient recombinant murine γ-herpesvirus-68. J. Immunol. 188, 1049–1056 (2012).

52. Crouse, J., Kalinke, U. & Oxenius, A. Regulation of antiviral T cell responses by type I interferons. Nat. Rev. Immunol. 15, 231–242 (2015).

53. Huber, J. P. & Farrar, J. D. Regulation of effector and memory T-cell functions by type I interferon. Immunology 132, 466–474 (2011).

54. Le Bon, A. et al. Type i interferons potently enhance humoral immunity and can promote isotype switching by stimulating dendritic cells in vivo. Immunity 14, 461–470 (2001).

55. Le Bon, A. & Tough, D. F. Links between innate and adaptive immunity via type I interferon. Current Opinion in Immunology 14, 432–436 (2002).

56. Aricò, E. et al. Humoral immune response and protection from viral infection in mice vaccinated with inactivated MHV-68: effects of type I interferon. J. Interferon Cytokine Res. 22, 1081–1088 (2002).

57. Bracci, L., La Sorsa, V., Belardelli, F. & Proietti, E. Type I interferons as vaccine adjuvants against infectious diseases and cancer. Expert Review of Vaccines 7, 373–381 (2008).

58. Proietti, E. et al. Type I IFN as a natural adjuvant for a protective immune response: lessons from the influenza vaccine model. J. Immunol. 169, 375–383 (2002).

59. Santini, S. M. et al. Type I interferon as a powerful adjuvant for monocyte-derived dendritic cell development and activity in vitro and in Hu-PBL-SCID mice. J. Exp. Med. 191, 1777–1788 (2000).

60. Cull, V. S., Broomfield, S., Bartlett, E. J., Brekalo, N. L. & James, C. M. Coimmunisation with type I IFN genes enhances protective immunity against cytomegalovirus and myocarditis in gB DNA-vaccinated mice. Gene Ther. 9, 1369–1378 (2002).

61. Bevan, M. J. Helping the CD8(+) T-cell response. Nat. Rev. Immunol. 4, 595–602 (2004).

62. MacLeod, M. K. L., Clambey, E. T., Kappler, J. W. & Marrack, P. CD4 memory T cells: what are they and what can they do? Semin. Immunol. 21, 53–61 (2009).

63. MacLeod, M. K. L., Kappler, J. W. & Marrack, P. Memory CD4 T cells: generation, reactivation and re-assignment. Immunology 130, 10–15 (2010).

64. Hwang, M. L., Lukens, J. R. & Bullock, T. N. J. Cognate memory CD4+ T cells generated with dendritic cell priming influence the expansion, trafficking, and differentiation of secondary CD8+ T cells and enhance tumor control. J. Immunol. 179, 5829–5838 (2007).

65. Gao, F. G. et al. Antigen-specific CD4+ T-cell help is required to activate a memory CD8+ T cell to a fully functional tumor killer cell. Cancer Res. 62, 6438–6441 (2002).

66. Blachère, N. E. et al. IL-2 is required for the activation of memory CD8+ T cells via antigen cross-presentation. J. Immunol. 176, 7288–7300 (2006).

67. Fitzgerald-Bocarsly, P., Dai, J. & Singh, S. Plasmacytoid dendritic cells and type I IFN: 50 years of convergent history. Cytokine Growth Factor Rev. 19, 3–19 (2008).

68. Kadowaki, N., Antonenko, S., Lau, J. Y.-N. & Liu, Y.-J. Natural Interferon α/β–Producing Cells Link Innate and Adaptive Immunity. The Journal of Experimental Medicine 192, 219–226 (2000).

69. McKenna, K., Beignon, A.-S. & Bhardwaj, N. Plasmacytoid dendritic cells: linking innate and adaptive immunity. J. Virol. 79, 17–27 (2005).

70. Karrich, J. J., Jachimowski, L. C. M., Uittenbogaart, C. H. & Blom, B. The Plasmacytoid Dendritic Cell as the Swiss Army Knife of the Immune System: Molecular Regulation of Its Multifaceted Functions. The Journal of Immunology 193, 5772–5778 (2014).

71. Obar, J. J., Fuse, S., Leung, E. K., Bellfy, S. C. & Usherwood, E. J. Gammaherpesvirus Persistence Alters Key CD8 T-Cell Memory Characteristics and Enhances Antiviral Protection. Journal of Virology 80, 8303–8315 (2006).

72. Badovinac, V. P. & Harty, J. T. Manipulating the rate of memory CD8+ T cell generation after acute infection. J. Immunol. 179, 53–63 (2007).

73. Doherty, P. C., Christensen, J. P., Belz, G. T., Stevenson, P. G. & Sangster, M. Y. Dissecting the host response to a gamma-herpesvirus. Philos. Trans. R. Soc. Lond., B, Biol. Sci. 356, 581–593 (2001).

74. Barton, E., Mandal, P. & Speck, S. H. Pathogenesis and host control of gammaherpesviruses: lessons from the mouse. Annu. Rev. Immunol. 29, 351–397 (2011).

75. H Speck, S. & W Virgin, H. Host and viral genetics of chronic infection: a mouse model of gamma-herpesvirus pathogenesis. Current Opinion in Microbiology 2, 403–409 (1999).

76. Virgin, H. W. & Speck, S. H. Unraveling immunity to gamma-herpesviruses: a new model for understanding the role of immunity in chronic virus infection. Curr. Opin. Immunol. 11, 371–379 (1999).

77. Simas, J. P. & Efstathiou, S. Murine gammaherpesvirus 68: a model for the study of gammaherpesvirus pathogenesis. Trends Microbiol. 6, 276–282 (1998).

78. Nash, A. A., Dutia, B. M., Stewart, J. P. & Davison, A. J. Natural history of murine -herpesvirus infection. Philosophical Transactions of the Royal Society B: Biological Sciences 356, 569–579 (2001).

79. Wu, T.-T., Blackman, M. A. & Sun, R. Prospects of a novel vaccination strategy for human gamma-herpesviruses. Immunol. Res. 48, 122–146 (2010).

80. Hwang, S. et al. Conserved herpesviral kinase promotes viral persistence by inhibiting the IRF-3-mediated type I interferon response. Cell Host Microbe 5, 166–178 (2009).

